# Hormonal and modality specific effects on males’ emotion recognition ability

**DOI:** 10.1101/791376

**Authors:** Adi Lausen, Christina Broering, Lars Penke, Annekathrin Schacht

## Abstract

Successful emotion recognition is a key component of our socio-emotional communication skills. However, little is known about the factors impacting males’ accuracy in emotion recognition tasks. This pre-registered study examined potential candidates, focusing on the modality of stimulus presentation, emotion category, and individual hormone levels. We obtained accuracy and reaction time scores from 312 males who categorized voice, face and voice-face stimuli for nonverbal emotional content. Results showed that recognition accuracy was significantly higher in the audio-visual than in the auditory or visual modality. While no significant association was found for testosterone and cortisol alone, the effect of the interaction with recognition accuracy and reaction time was significant, but small. Our results establish that audio-visual congruent stimuli enhance recognition accuracy and provide novel empirical support by showing that the interaction of testosterone and cortisol modulate to some extent males’ accuracy and response times in emotion recognition tasks.

## Introduction

Emotion recognition is a basic skill thought to carry clear advantages for predicting behaviour, as well as forming and maintaining social bonds (Soto & Levenson, 2009). Intriguingly, research on sex differences highlights that males are less accurate than females when completing emotion recognition tasks (e.g., Thompson & Voyer, 2014; Hall, 1984). However, effect sizes are comparably small and multiple factors known to impact the ability to recognize emotions have yet to be fully controlled for (e.g., Hall et al., 2000; see Chaplin, 2015; Fischer, & LaFrance, 2015; Hyde, 2014; Schirmer, 2013, for an overview regarding explanations for sex-based behaviour patterns). The ability to correctly interpret emotional expressions forms the basis of social interactions and personal relationships (e.g., Fischer & Manstead, 2008; Keltner & Kring, 1998) yet, there is a lack of direct evidence for reasons why males have an often assumed disadvantage when it comes to accurately recognizing emotions. Therefore, the main aim of this study was to systematically investigate potential factors that might impact males’ ability to recognize emotions.

One of the factors supposed to impact emotion recognition is the modality of stimulus presentation (Hall, 1984). In many everyday situations, judgments about others’ emotional states require the integration of information from various sensory modalities making use of different cues such as facial expressions, tone of voice (i.e., prosody), or body language (Klasen et al., 2014). Thus, it has been argued that emotion recognition is a multimodal event (Piwek et al., 2015). Indeed, a growing number of studies have pointed out that in emotion recognition tasks the stimuli presented in isolation (i.e., visual or auditory) have lower accuracy scores and slower response times than the audio-visual presentation of emotional expressions (Jessen et al., 2012; Paulmann & Pell, 2011; Baenziger et al., 2009; Collignon et al., 2008; Kreifelts et al., 2007; de Gelder & Vroomen, 2000). Research on unimodal emotion recognition reported that emotions are better recognized from faces than from voices (e.g., Waaramaa, 2017).

However, these observations were often contradictory (e.g., Kraus, 2017). Furthermore, previous research in the unimodal domains highlighted that specific emotions are not recognized equally well in the auditory and visual modality. In studies on the vocal channel, participants were faster and most accurate to recognize anger (e.g., Chronaki et al., 2018; Cornew et al., 2009; Juslin & Laukka, 2003), while in studies on facial expressions, happiness was shown to be recognized more accurately and faster than any other emotion (e.g., Kosonogov & Titova, 2018; Wells et al., 2016; Nummenmaa & Calvo, 2015; Williams et al., 2009; Montagne et al., 2007; Palermo & Coltheart, 2004; Elfenbein & Ambady, 2002). Despite these converging patterns, it is as yet not possible to make definite claims regarding the advantage of certain emotional categories because, at least within the vocal domain, recognition accuracy (RA) was found to be strongly influenced by the type of stimulus used (see Lausen et al., 2019, for an overview). Whether the voice is a more reliable source than the face in emotion recognition tasks has been rarely pursued, and results are limited to specific emotions, paradigms, as well as, by a number of methodological differences between studies. Thus, until further evidence regarding RA within specific sensory modalities and emotional categories is provided, the direction of these effects remains an open question.

A recently emphasized influence on the ability to recognize emotions concerns potential effects of steroid hormones, such as testosterone (Gignell et al., 2019). Testosterone (T) receptors are distributed throughout the nervous system with high concentrations in areas associated with emotional processing such as the hypothalamus and amygdala [see Gignell et al., 2019, for details]. However, only few studies have assessed the influence of T concentrations on emotion recognition in both sexes and an even smaller subsection has specifically addressed the impact of T levels on males’ ability to recognize emotions. For example, an fMRI study by Derntl et al. (2009) investigated the influence of blood T levels on males’ RA in an explicit emotion recognition task. Results showed increased amygdala activity in individuals with high T levels during the presentation of fearful and angry faces. In addition, the authors found that reaction times (RTs) to fearful male faces negatively correlated with T level concentrations. However, no correlation was found between RA and T levels. Subsequent studies reported a negative correlation between salivary T levels and emotion recognition in male adolescent groups (Fujisawa & Shinohara, 2011) or found a positive correlation between higher levels of T and emotion recognition (Vongas & Al Hajj, 2017). By presenting participants with emotional facial expressions at two different intensity levels (i.e., 50% and 100%), Rukavina et al. (2018) found that RA decreases when salivary T is high, especially for full-blown expressions of sadness and for disgust when presented at 50% intensity. Based on these findings, the authors concluded that RA decreases with increasing levels of T.

These contradictory findings are likely the result of a number of methodological differences such as insufficient statistical power (i.e., sample sizes ranging from 21 to 84 males), T assessment from blood or saliva, as well as storage and analyses of hormone samples (see Schultheiss et al., 2019, for details). Another possible explanation for the discrepancies is that another hormone, cortisol (C), may constrain T influence on emotion recognition. C, an end product of the hypothalamic-pituitary-adrenal (HPA) axis, was found to inhibit T by reducing hypothalamic-pituitary-gonadal (HPG) activity and blocking androgen receptors [see Sarkar et al., 2019; Viau, 2002, for details]. To reconcile mixed findings on the roles of T and C in human social behavior, Mehta and Josephs (2010) proposed the dual-hormone hypothesis. According to this hypothesis T predicts a wide range of behaviors, but only under the condition that C concentrations are low. If C concentrations are high, the T-behavior association is supposed to be attenuated (Mehta & Prasad, 2015; Carré & Mehta, 2011). This hypothesis was supported in a variety of studies, which demonstrated that across different psychological domains the interaction between T and C influences empathy, as well as, dominant, status-relevant, risk-taking and antisocial behavior (see Sarkar et al., 2019, for an overview). However, it should be noted that other studies report only small effects (e.g., Dekkers et al., 2019; Grebe et al., 2019) null-findings [e.g., Mazur & Booth, 2014], and even reversed patterns [i.e., T was related to status-relevant behavior or facial dominance for high but not low C (e.g., Kordsmeyer et al., 2018; Welker et al., 2014)] for the dual-hormone hypothesis. Considering the interaction between the HPG and HPA axes might nevertheless lead to more reliable predictions regarding emotion recognition than the assumption of a single-hormone association (Sarkar et al., 2019; Carré & Mehta, 2011).

Based on the above-mentioned findings, the present study had three major aims. Firstly, it aimed at examining whether males’ RA is influenced by the modality of stimulus presentation. We hypothesized that RA would be better in the audio-visual modality than in the auditory or visual modality (1a), and lower in the visual compared to the auditory modality (1b). Second, we aimed to replicate previous findings by examining the extent RA and RTs vary across discrete emotion categories as a function of modality (e.g., Lambrecht et al., 2014). Specifically, we expected higher accuracy scores and faster RTs for disgusted, fearful and sad expressions in the audio-visual than in both the auditory and the visual modality (2a). We also hypothesized that angry expressions would be identified faster and with higher accuracy in the vocal compared to the facial domain, while for happy expressions we expected the reverse pattern (2b). A third aim was to alleviate some of the methodological flaws of previous research by using a large sample size to examine whether variations in males’ ability to recognize emotions are due to T level concentrations. We expected a negative correlation between T and RA (3a), and that participants with high levels of T would specifically react faster to angry and fearful expressions (3b)^1^. In addition, we conducted an exploratory analysis on the associations between C and RA, C and RT, as well as on the relationship between RA or RT and the interaction between T and C levels.

## Method

The study was approved by the ethics committee of the Georg-Elias-Mueller-Institute of Psychology (University of Goettingen), and conducted in accordance with the ethical principles formulated in the Declaration of Helsinki (2013). Participants gave informed consent and were reimbursed with course credit or 8 Euros per hour.

### Participants

A total of 312 males (age range 18-36 years; *M*_*Age*_ = 24.3, *SD* = 3.7) were recruited on the university campus using flyers and the Institute of Psychology participant database (*ORSEE*, www.orsee.org), as well as by posts on the social media site Facebook and the online platform Stellenwerk Jobportal University Goettingen (www.stellenwerk-goettingen.de). Of the 312 recruited subjects, 30 participants were excluded from analysis due to self-reported hearing problems, psychiatric or neurological disorders, or intake of psychotropic/hormone medication. After these exclusions, a total of 282 participants with a mean age of 24.3 years (*SD* = 3.8) were included in the analysis.

#### Stimulus material

Stimuli were displayed under three experimental modality conditions: auditory, visual and audio-visual. In each experimental condition, stimuli were presented in one of the emotions of interest (i.e., anger, disgust, fear, happiness, sadness) as well as in a neutral state (i.e., baseline expression).

#### Audio stimuli

The audio stimuli consisted of pseudo-speech (i.e., pseudo-words, pseudo-sentences) and non-verbal vocalizations (i.e., affect bursts). We decided to use pseudo-speech (i.e., a language devoid of meaning) and non-verbal vocalizations as they have been argued to capture the pure effects of emotional prosody independent of lexical-semantic cues and, to be an ideal tool when investigating the expression of emotional information when there is no concurrent verbal information present (Pell et al., 2015; Banse & Scherer, 1996). The stimuli were sampled from well-established databases or provided by researchers who developed their own stimulus materials. We validated all stimuli in a previous study (cf. Lausen et al., 2019; Lausen & Schacht, 2018) and selected only a subset of stimuli (i.e., with the highest accuracy) from each database (see Table 1).

**Table 1.** Description of audio materials

| Database | Speakers | Emotions | Nature of material | Number of stimuli selected | Total stimuli |
| --- | --- | --- | --- | --- | --- |
| *Magdeburg Prosody Corpus*<br>(Wendt & Scheich, 2002) | 2 actors<br>(1 male/1 female) | Anger, disgust,<br>fear, happiness,<br>sadness, neutral | Pseudo-words | 4 | 48 |
| *Paulmann Prosodic Stimuli*<br>(Paulmann & Kotz, 2008;<br>Paulmann et al., 2008) | 2 actors<br>(1 male/1 female) |  | Pseudo-sentences | 4 | 48 |
| *Montreal Affective Voices*<br>(Belin et al., 2008) | 8 actors<br>(4 male/4 female) |  | Affect bursts |  | 48 |

The physical volume of stimulus presentations across the nine laptops used in the experiment was controlled by measuring sound volume of the practice trials with a professional sound level meter, *Nor140* (Norsonic, 2010, Lierskogen, Norway). No significant difference in volume intensity was observed [*F*_(8,40)_ = 1.546, *p* = 0.173].

#### Visual stimuli

Visual stimuli consisted of 24 frontal face photographs (12 males/12 females) extracted from the *Radboud Faces Database* (Langner et al., 2010). The presentation time of the faces was matched to the length of the voice stimuli (i.e., from 319 ms to 4821 ms). A gray ellipsoid mask, ensuring a uniform figure/ground contrast surrounded the stimuli, with only the internal area of the face visible (9×14 cm, width and height). The stimuli were presented in colour and corrected for luminance across emotion conditions [*F*_(5,137)_ = 0.200, *p* = 0.962], using *Adobe Photoshop CS6* (Version 13.0.1, 2012, San Jose, CA).

#### Audio-visual stimuli

The voice stimuli were simultaneously presented with the face stimuli. Using *Adobe Premiere Pro CS6* (Version 6.0.5) videos were created, matching face and voice stimuli for sex and emotion category.

### Procedure, experimental task and saliva samples

Participants were informed that the study required them to provide two saliva samples over a period of about two hours. A day before the main experiment, they were sent an email instructing them to abstain from sports and the consumption of alcohol, drugs or unnecessary medication on the day of the study. Furthermore, they were instructed not to consume drinks containing caffeine within three hours of the experiment and to refrain from eating, drinking (except water), smoking and brushing their teeth within one hour of the experiment. Adherence to these instructions was assessed using a screening questionnaire (Schultheiss & Stanton, 2009). As individual differences in peak hormone levels measured in the morning have been argued to be a better predictor of behavioural responses to emotional stimuli than measurements later in the day (Schultheiss & Stanton, 2009), the designated time slot for testing was between 9:00am to 11:00am.

Participants were tested in groups of up to nine individuals. On the day of the study, after completing the consent form, participants received oral and written instructions about the procedure of the experiment and the collection of saliva samples. The saliva samples were collected before (T1) and after (T2) the Emotion Recognition Task^2^. The experiment was programmed using *Python* (Version 2.7.0, Python Software Foundation, Beaverton, OR) and run on a *Dell Latitude E5530 Laptop* with a 15.6 LCD display screen. The audio stimuli were presented binaurally via headphones (*Bayerdynamic DT 770 PRO*).

#### Emotion recognition task

The emotion recognition task consisted of three blocks, each block displaying one of the three experimental conditions: auditory, visual, and audio-visual. Each experimental condition contained 144 stimuli. A permutation was applied to randomize the order in which the experimental conditions were presented to the participants. Six different permutations were created, and each permutation was allocated randomly in blocks of six participants. The order of the stimuli within each experimental condition was completely randomized. The audio and visual stimuli were matched for duration, sex, and emotion category (see *Table S1* in supplementary material for an example of how the audio and visual stimuli were matched). Before each experimental condition, participants were familiarized with the task in a short training session comprised of three stimuli. Each trial began with a blank screen followed by a fixation cross. Following the presentation of a stimulus, a circular answer display appeared, containing all six categories of interest (i.e., anger, disgust, fear, happiness, sadness, neutral) and the selection cursor, which appeared in the centre of the display. The sequence of the emotion labels was randomized for each participant and remained the same throughout the task. Participants had to select an emotion category, using the mouse to move the cursor, before the next stimulus was presented. Reaction times were measured, starting with the onset of the answer display and ending with the participant’s response. Figure 1 displays the time course of the emotion recognition task.

**Figure 1.**
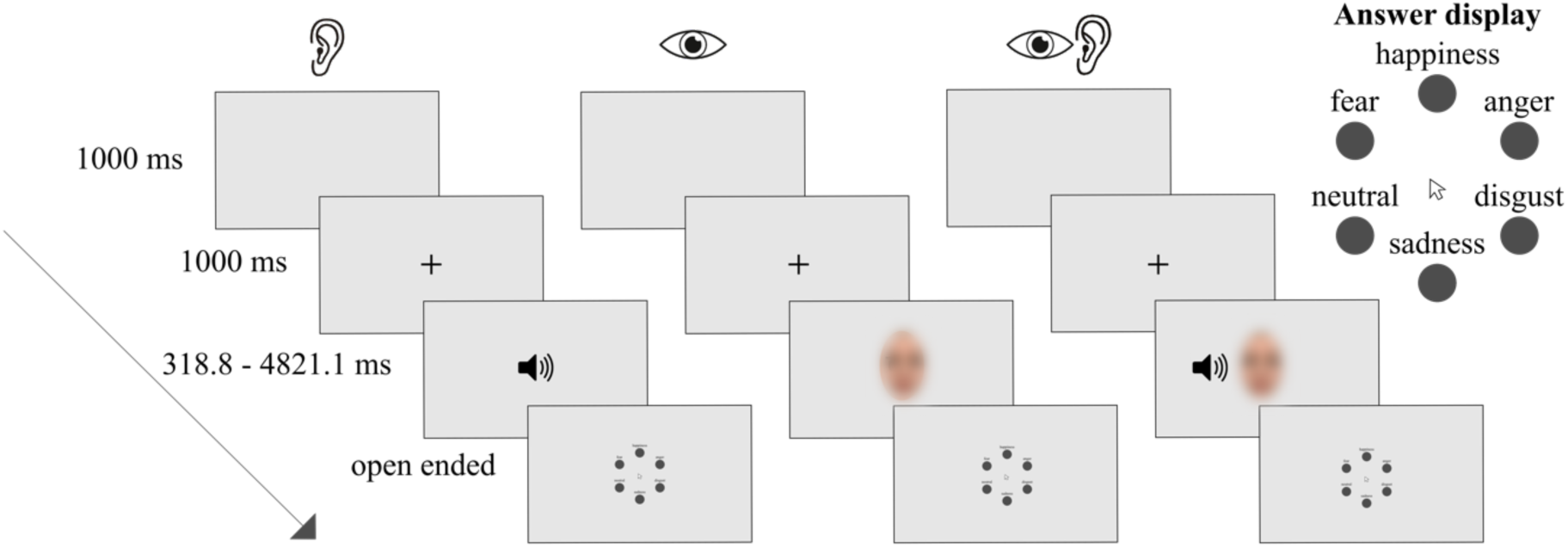
Emotion recognition task. Each trial began with a blank screen (shown for 1000ms) which was followed by a fixation-cross appearing at the center of the screen (for 1000ms) at which participants were asked to fixate throughout the trial. After the presentation of the stimulus a circular answer display containing all six categories of interest (i.e., anger, disgust, fear, happiness, sadness, neutral) and the selection cursor (which appeared in the center of the display) were presented. The responses were made by using the mouse to move the cursor. Reaction times were measured, starting with the onset of the answer display and ending with the participant's response. There was no time limit for emotion judgments. Participants could hear/see the stimulus only once. The presentation of the stimuli was initiated by pressing the Spacebar-key at the beginning of each block. At the end of each block a visual message in the center of the screen instructed participants to take a break if they wished to or to press the Spacebar-key to proceed with the next block. (The face stimuli were obscured as per bioRxiv policy)

#### Saliva samples

The two saliva samples (2 ml per sample) were collected from each participant via passive drool through a straw (Schultheiss et al., 2012) into an *IBL SaliCap* sampling device. These plastic vials were stored frozen at −80°C until shipment on dry ice to the Endocrinology Laboratory at Technical University of Dresden. At this facility, the samples were analysed for T and C levels via chemiluminescence immunoassays with high sensitivity (IBL International, Hamburg, Germany). The intra- and inter-assay coefficients of variation for T were < 11% and for C < 8%. For T the variance between participants was 14.81% and 3.85% within participants with an intra-class correlation coefficient (ICC) of 79.35%, while for C the variance between participants was 23.78% and 28.20% within participants with an ICC of 45.74%. As the distributions of T and C were positively skewed (T_skewness_ = 1.56; C_skewness_ = 1.49) a log-transformation was performed (e.g., Mehta et al., 2015). The log-transformation reduced skewness substantially [log(T) skewness = −0.06; log(C) skewness = 0.01]. Outliers were winsorized to ± 3 standard deviations (Mehta et al., 2015).

### Study design and power analysis

A balanced within-subjects factorial design was fitted to assess males’ judgments of emotions. The design was balanced for modalities, emotion categories and encoder sex in each stimulus type. Independent within-participant factors were modalities, emotion categories, stimuli types, and encoder sex. Independent between-participant variables were T and C. Dependent variables were RA and RT.

A target sample size of 231 males was determined using an approximate correlation power analysis, Bonferroni-corrected for multiple testing (*r* = .25; *α* = .05/20; *1 – β* = .80). To account for possible attrition, the sample size was increased by a minimum of 14%.

### Statistical analysis

In line with our preregistration, the primary analysis for our first and second hypotheses was performed using *Friedman*- and *Wilcoxon-rank-sum* tests. For the association between the dependent variables (RA, RT) and T levels we ran *Spearman* correlations (H3a, b). The exploratory analyses of the quantitative variables T and C were performed using generalized linear models (*quasi-binomial logistic regression*) for the binary response variable emotion recognition and linear models for the response variable reaction time, which was normalized by log transformation. To obtain a more reliable value and to cover the observation interval, the two baseline measures for T and C were averaged (Kordsmeyer et al., 2018; Idris et al., 2017). The dispersion parameter of the quasi-binomial model accounted for dependencies caused by repeated measurements within the participants. Modality and emotion category were fitted as nominal variables and stimulus duration as quantitative variable. The interaction of the quantitative variables T and C was fitted by the product of both variables as an additional predictor. Tertiles for both variables, T and C, were fitted to investigate more general interaction patterns and to reduce the influence of T and C extreme values on the model equation. Chi-square tests of the deviance analysis and F-tests of the analysis of variance were used to analyse effects of predictor variables. In the quasi-binomial logistic regression, odds ratio (OR) were used to compare emotion recognition accuracies. RTs were compared by the difference of the means. Tukey’s method of multiple pairwise comparisons was used to compute simultaneous 95% confidence intervals for both, OR and mean differences.

For the descriptive analysis of the data, *relative frequencies*, *confusion matrices* and *Wagner’s* (1993) *unbiased hit rate* (*H*_*u*_), which is the rate of correctly identified stimuli multiplied by the rate of correct judgments of the stimuli, were calculated. The data was analysed using the R language and environment for statistical computing and graphics version 3.4.3 (R Core Team, 2017) and the integrated environment R-Studio version 1.0.153 (used packages: *pwr*; *MASS*; *coin*; *glm*; *multcomp*; *mvtnorm*; *ggplot2*).

## Results

### Descriptive analysis

Audio-visual emotional expressions were recognized with approximately 90% accuracy (lowest identification rate 89% for disgust). Angry expressions were recognized with better accuracy from the voice (90%) than the face (82%). Conversely, for fearful, happy and sad expressions accuracy scores were higher when presented visually (85% ≤ *accuracy scores* ≤ 99%) than auditorily (72% ≤ *accuracy scores* ≤ 77%). Neutral expressions had high accuracy scores in all three conditions of stimulus presentation (90% ≤ *accuracy scores* ≤ 95%). Participants were faster at recognizing disgust, fear, happy, sad and neutral expressions in the visual and audio-visual modalities (median (*Md*) values between 1.03 sec. to 1.46 sec.) than in the auditory modality (*Md* values between 1.50 sec. to 1.95 sec.). Although the RTs for disgusted, sad and neutral expressions were similar in the visual and audio-visual modalities, participants were slightly faster at recognizing fear and happy in the visual than audio-visual modality. For angry expressions, the RTs were much shorter in the audio-visual (1.23 sec.) than in the auditory and visual modality, but much longer in the visual (1.53 sec.) than in the auditory modality (1.47 sec.). Figure 2 illustrates participants’ RA (panel A) and RTs (panel B) by modality and emotion categories.

**Figure 2.**
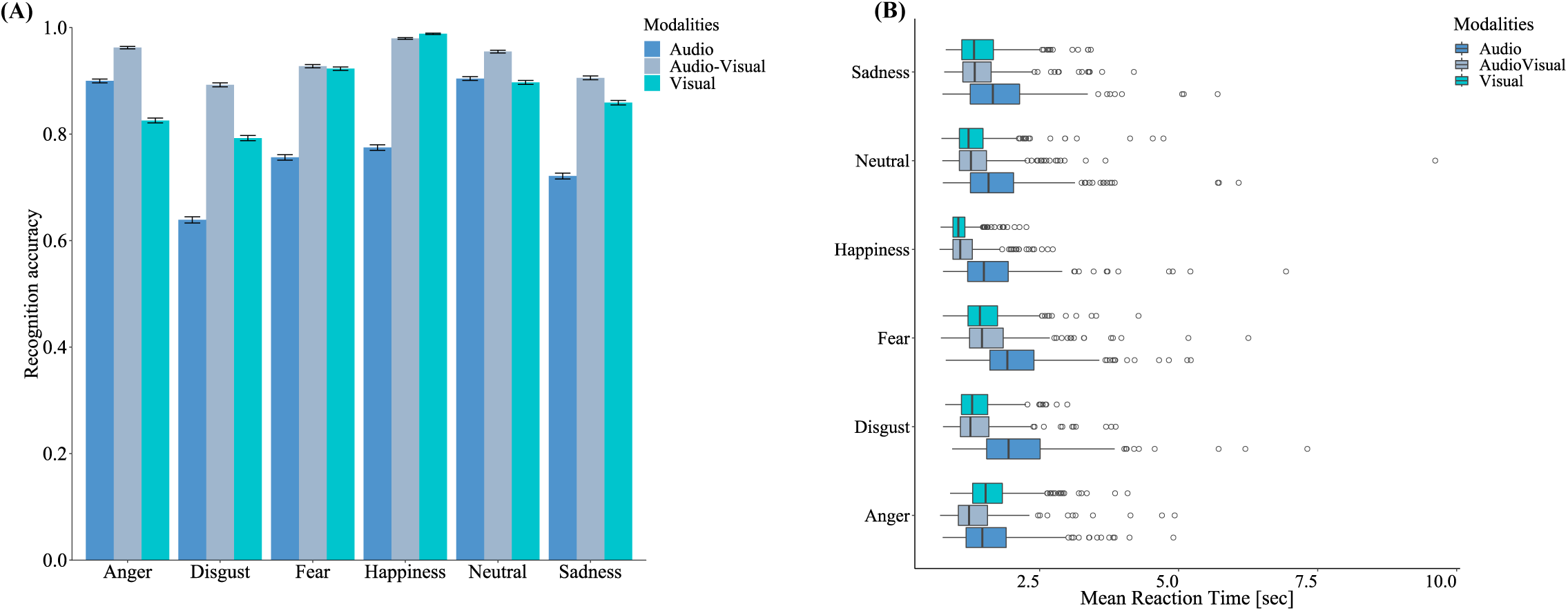
Recognition accuracy (RA) and reaction times (RTs) by modality and emotion categories. The bar charts (**panel A**) display RA, while the boxplots (**panel B**) illustrate the mean RT distributions. Error bars represent the standard error. The boxplots indicate that the distributions of RT are right skewed.

In all three modalities participants often misclassified happy and sad expressions as neutral. In the auditory and audio-visual modalities angry was mistaken for fearful, neutral for angry and fearful for sad. In the visual modality fear was confused with disgust, whereas anger and neutral were confused with sadness. Participants frequently misclassified disgust with anger in the visual and audio-visual modalities, while in the auditory modality disgust was mistaken for neutral. The error classification patterns along with the unbiased hit rates are presented in Table 2.

**Table 2.** Confusion Matrices and unbiased hit rates (*H*_*u*_) for participants judgments of emotion categories

| Modality | Emotions portrayed | Emotion judgments | | | | | | | $H_u$ |
| --- | --- | --- | --- | --- | --- | --- | --- | --- | --- |
|  |  | Anger | Disgust | Fear | Happiness | Neutral | Sadness | Total |  |
| Auditory | Anger | **6089** | 59 | 267 | 152 | 175 | 26 | 6768 | .766 |
|  | Disgust | 347 | **4324** | 438 | 280 | 815 | 564 | 6768 | .590 |
|  | Fear | 162 | 173 | **5118** | 96 | 406 | 813 | 6768 | .621 |
|  | Happiness | 116 | 27 | 15 | **5243** | 1335 | 32 | 6768 | .665 |
|  | Neutral | 339 | 52 | 62 | 159 | **6119** | 37 | 6768 | .549 |
|  | Sadness | 97 | 50 | 335 | 175 | 1230 | **4881** | 6768 | .554 |
|  | Total | 7150 | 4685 | 6235 | 6105 | 10080 | 6353 | 40608 | — |
| Visual | Anger | **5587** | 244 | 194 | 6 | 234 | 503 | 6768 | .638 |
|  | Disgust | 1288 | **5363** | 48 | 13 | 41 | 15 | 6768 | .704 |
|  | Fear | 51 | 282 | **6245** | 14 | 73 | 103 | 6768 | .847 |
|  | Happiness | 6 | 2 | 11 | **6689** | 59 | 1 | 6768 | .967 |
|  | Neutral | 167 | 15 | 47 | 102 | **6071** | 365 | 6767* | .791 |
|  | Sadness | 135 | 134 | 262 | 11 | 412 | **5814** | 6768 | .734 |
|  | Total | 7234 | 6040 | 6807 | 6835 | 6890 | 6801 | 40607 | — |
| Audio-visual | Anger | **6513** | 46 | 91 | 8 | 71 | 39 | 6768 | .860 |
|  | Disgust | 505 | **6040** | 69 | 14 | 81 | 59 | 6768 | .858 |
|  | Fear | 39 | 155 | **6277** | 9 | 92 | 196 | 6768 | .873 |
|  | Happiness | 5 | 2 | 7 | **6629** | 121 | 4 | 6768 | .969 |
|  | Neutral | 170 | 11 | 25 | 35 | **6462** | 65 | 6768 | .859 |
|  | Sadness | 55 | 27 | 196 | 9 | 353 | **6128** | 6768 | .855 |
|  | Total | 7287 | 6281 | 6665 | 6704 | 7180 | 6491 | 40608 | — |
| Across all 3 modalities | Anger | **18189** | 349 | 552 | 166 | 480 | 568 | 20304 | .752 |
|  | Disgust | 2140 | **15727** | 555 | 307 | 937 | 638 | 20304 | .716 |
|  | Fear | 252 | 610 | **17640** | 119 | 571 | 1112 | 20304 | .780 |
|  | Happiness | 127 | 31 | 33 | **18561** | 1515 | 37 | 20304 | .864 |
|  | Neutral | 676 | 78 | 134 | 296 | **18652** | 467 | 20303* | .710 |
|  | Sadness | 287 | 211 | 793 | 195 | 1995 | **16823** | 20304 | .709 |
|  | Total | 21671 | 17006 | 19707 | 19644 | 24150 | 19645 | 121823 | — |

Note: Frequencies of correctly judged portrayals are given on the main diagonal in boldface type. \*If the number is less than the planned number of emotion judgments that is due to recording failure. $H_u$ = the rate of correctly identified stimuli multiplied by the rate of correct judgments of the stimuli.

### Main analysis

#### *Recognition accuracy in the three modalities* [Aim 1]

Participants’ RA was significantly influenced by the modality of stimulus presentation (Friedman test: *χ^2^*_(2)_ = 448.56, p < 0.001). The results of Wilcoxon-rank-sum test indicated that RA was significantly higher in the audio-visual modality than in the visual (*z* = 12.99, *p* < 0.001, _*95%*_*CI* = [0.052; 0.062], effect size (*r*) = 0.774) or auditory modality (*z* = 14.525, *p* < 0.001, _*95%*_*CI* = [0.146; 0.163], *r* = 0.865). Participants’ were also significantly more accurate at discriminating emotions when making judgments on visual than on audio stimuli (*z* = 13.553, *p* < 0.001, _*95%*_*CI* = [0.090; 0.108], *r* = 0.807). Figure 3 illustrates RA in the three conditions of stimulus presentation.

**Figure 3.**
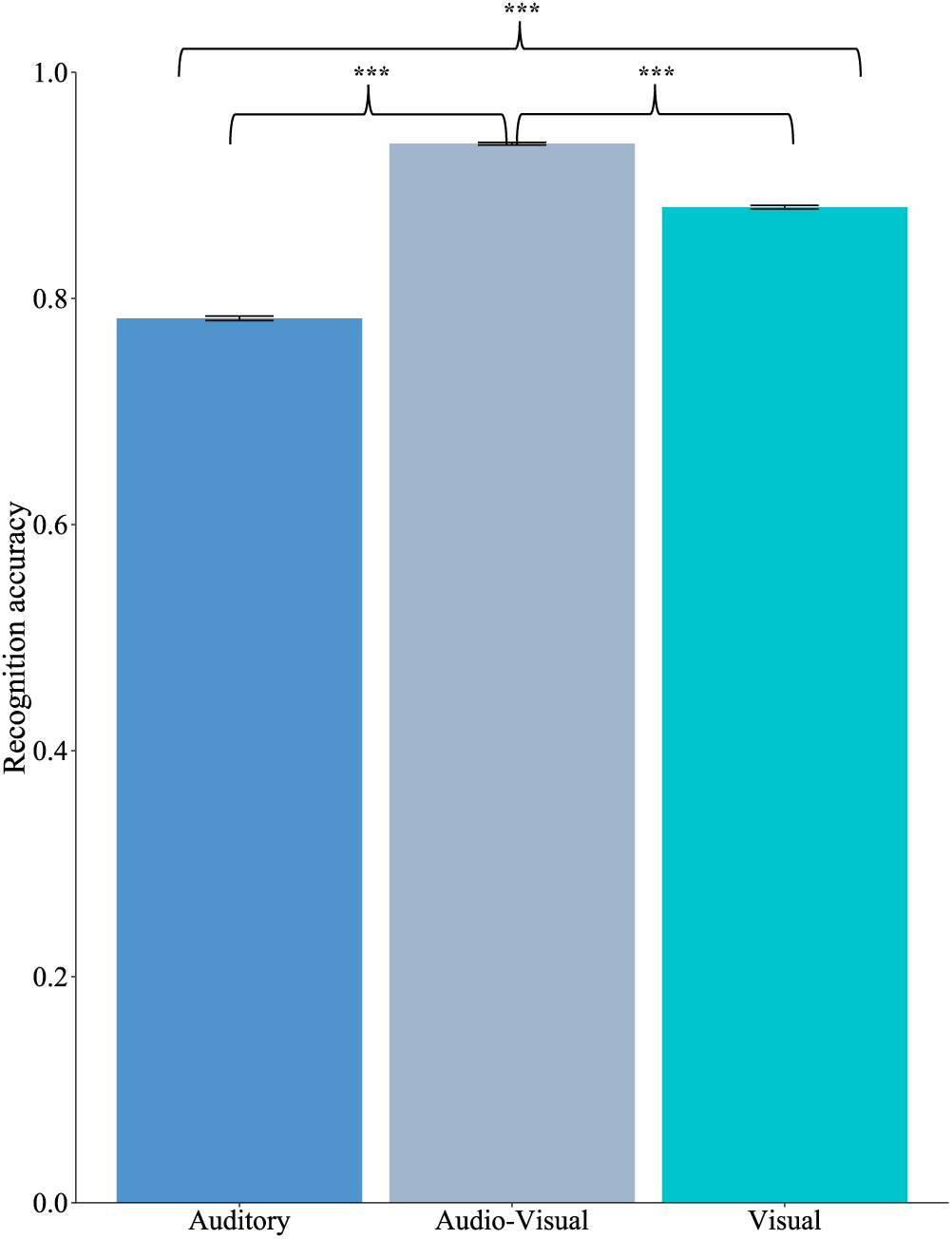
Bar chart showing the recognition accuracy (RA) in the three conditions of stimulus presentation. Error bars represent the standard error. RA was significantly higher for the audio-visual presented stimuli than for the visual-or auditory stimuli. Accuracy scores were significantly higher for the visual-than for auditory condition.

#### *Emotion specificity and modality* [Aim 2]

The modality of stimulus presentation across fearful, disgusted and sad expressions significantly influenced participants’ RA (Friedman test: *χ^2^*_(2)_ = 400.47, *p* < 0.001) and RTs (Friedman test: *χ^2^*_(2)_ = 208.77, *p* < 0.001). Results comparing RA and RTs between modalities for each emotion category showed that participants were significantly more accurate and faster at categorizing these emotions in the audio-visual than auditory modality (*p*s < 0.001; effect sizes for accuracy ranging from 0.813 < *r* < 0.852 and for RTs ranging from 0.422 < *r* < 0.760). Although RA was significantly higher for disgust (*p* < 0.001; *r* = 0.605) and sad expressions (*p* < 0.001; *r* = 0.417) in the audio-visual than visual modality, the accuracy scores for fear did not significantly differ between these two modalities (*p* = 1.00; *r* = 0.038). Similarly, we observed no significant RT differences between the audio-visual and visual modality for these three emotions (*p*s > 0.05; 0.005 < *r* < 0.159). While participants were significantly better at recognizing angry expressions in the voice than in the face (*p* < 0.001, *r* = 0.492), RTs did not differ significantly between these two modalities (*p* = 1.00, *r* = 0.052). In contrast, happy, disgusted, fearful, and sad expressions had significantly higher accuracy scores and faster RTs when they were presented visually than auditorily (*p*s < 0.001; 0.625 < *r*_Accuracy_< 0.868; 0.487 < *r*_RT_ < 0.816). Table 3 displays the test statistics for each modality and emotion category.

**Table 3.**
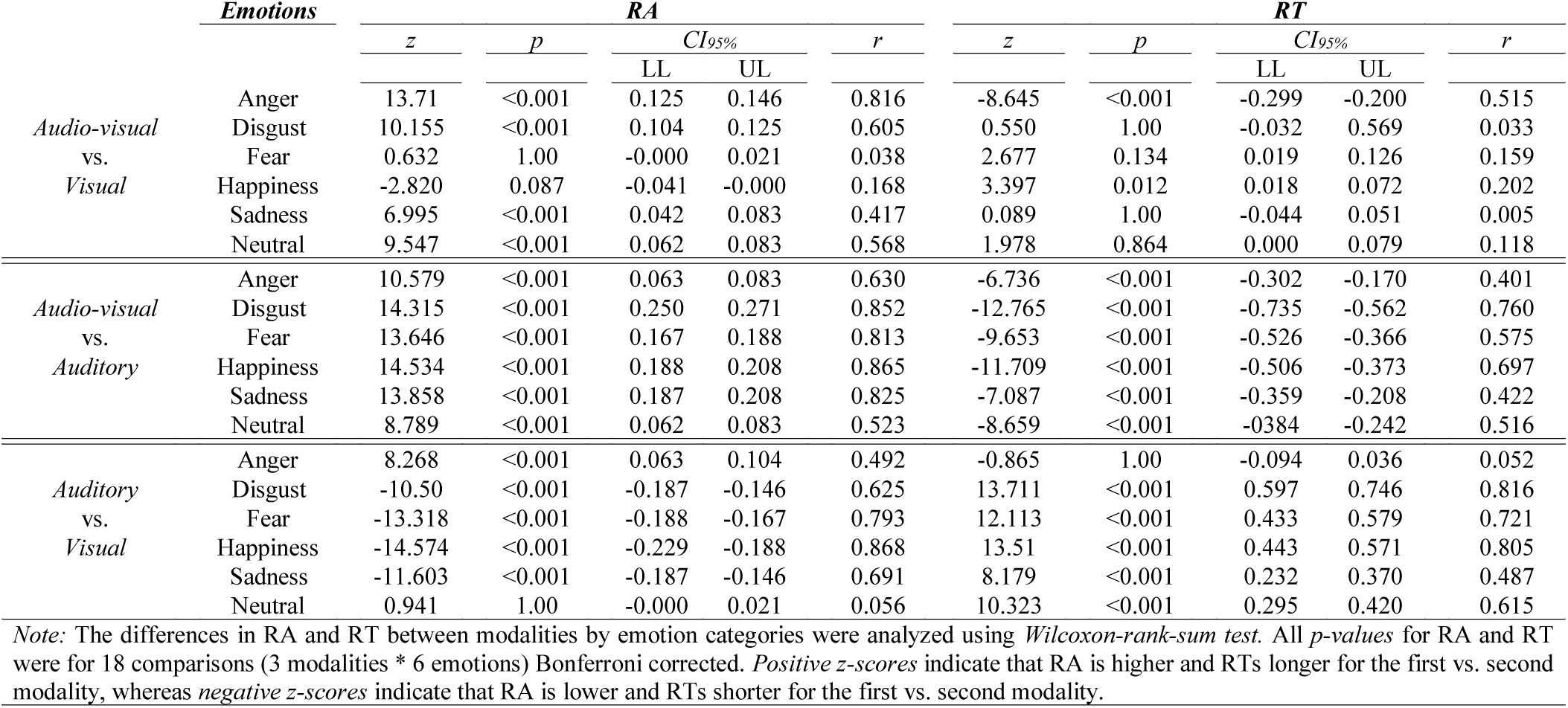
Recognition accuracy (RA) and reaction times (RTs) standardized *z-scores*, *p-values*, *95% confidence intervals* (CI_95%_) and *effect sizes* (*r*) for the comparisons between modalities by emotion categories

#### *Interplay of hormones, recognition accuracy and reaction times* [Aim 3]

Spearman’s rank correlation coefficient between T1 and T2 for T was *r*_*s*_ = 0.79 and *r*_*s*_ = 0.60 for C. No significant associations between T or C and RA/RTs were found (*p*s > .05; correlation coefficients (*r*_*s*_) close to zero; *Figure S1* in supplementary material illustrates the relationship between T or C and RA/RTs, also across all modalities). Similarly, there were no significant associations between T or C and RA/RTs for specific emotion categories (see *Table S2* in supplementary material). Logistic and linear models, however, showed that the interaction between testosterone and cortisol (TxC) significantly influenced participants’ RA (*χ^2^*_(4)_ = 46.30, *p* < 0.001, *r* = 0.022) and RTs (*F*_(4, 121806)_ = 8.26, *p* < 0.001, *r* = 0.016). *Table S3* in supplementary material provides an overview on the model terms and the corresponding statistics for both RA and RTs. The odds ratio estimates for RA and the linear contrasts for the pattern of the differences in RTs for all combinations between T and C terciles showed that participants RA was significantly higher for *T*_*High*_/*C*_*Low*_ and *T*_*Low*_/*C*_*High*_, but lower for *T*_*Middle*_/*C*_*Low*_ or *T*_*Low*_/*C*_*Middle*_. RTs were shorter for *T*_*High*_/*C*_*Low*_, *T*_*Low*_/*C*_*Low*_, as well as for *T*_*Low*_/*C*_*Middle*_. For the combinations *T*_*High*_/*C*_*High*_or *T*_*Middle*_/*C*_*High*_ RTs were significantly longer. In Figure 4, panels **A**, **B** display the corresponding statistics for all comparisons between T and C terciles, while panels **A**_**I**_, **B**_**I**_ illustrate the conditional patterns.

**Figure 4.**
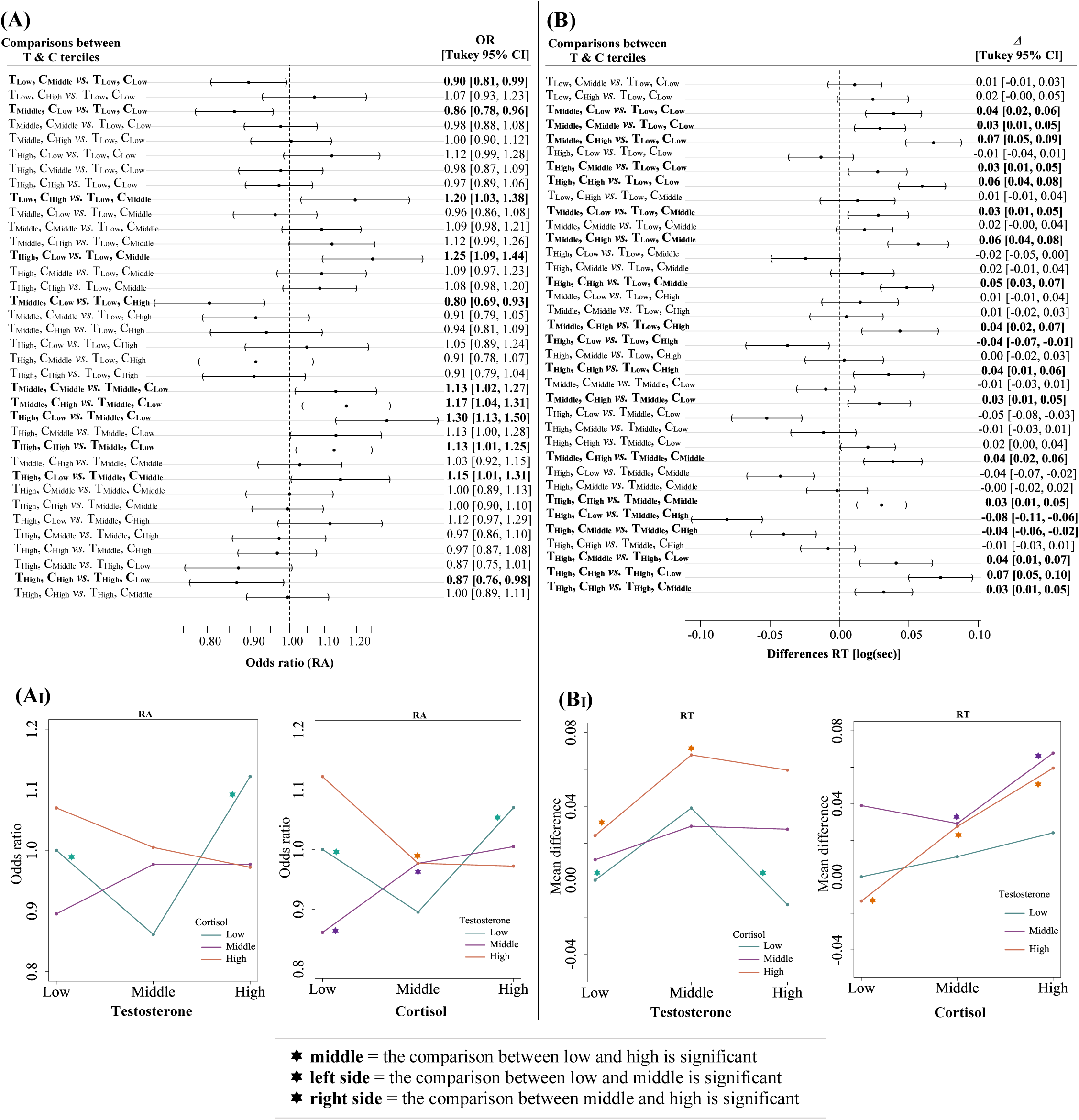
Pairwise comparisons and conditional patterns of T and C terciles combinations for recognition accuracy (RA) and tion time (RT) The comparisons between hormone terciles for RA are illustrated in panel **(A)**, while the linear contrasts for the pattern of the rences in RT are illustrated in panel **(B)**. The significant combinations are highlighted in bold. The T pattern conditional under d C pattern conditional under T for RA are shown in panel **(A_I_)** and panel **(B_I_)** for RT. In panel**(A)** odds ratio for combination 1 (e.g., *T*_*High*_/*C*_*High*_) vs. combination 2 (e.g., *T*_*High*_/*C*_*Low*_) less than 1 indicate that the recognition ability for combination 2 (*T*_*High*_/*C*_*Low*_) is higher than for combination 1 (*T*_*High*_/*C*_*High*_), whereas values greater than 1 vice-versa. If adds ratio of 1 is included in the confidence interval, the difference in the recognition probabilities is not significant. In panel **(B)** negative differences of RT for combination 1 (e.g., *T*_*High*_/*C*_*High*_) vs. combination 2 (e.g., *T*_*High*_/*C*_*Low*_) indicate that the RT for bination 2 (*T*_*High*_/*C*_*High*_) are longer than for combination 1 (*T*_*High*_/*C*_*Low*_), whereas positive differences vice-versa. If the difference ro is included in the 95%CI, the difference in RT is not significant. As it can be observed, for ***T conditional under C***_***Low***_ and ***C conditional under T***_***Low***_ there is a quadratic relationship [i.e., the accuracy eases from low to middle T or C and then increases from middle to high T or C (see panel A_I_); for ***T conditional under C***_***Low***_ the ncreases from low to middle T and then decreases from middle to high T (see panel B_I_)]. For ***C conditional under T***_***High***_ the ionship is monotone [i.e., the accuracy decreases from low C to high C (see panel A_I_); the RT increases from low C to high C panel B_I_)].

## Discussion

The main objective of the present study was to investigate whether males’ RA is influenced by the modality of stimulus presentation in an explicit emotion recognition task. In addition, we examined whether specific emotions are more quickly and accurately detected as a function of modality. Finally, we explored the effects of individual differences in T and C, as well as their interaction with RA and RTs. Our results provide compelling evidence that RA is greatly improved when visual and audio information were jointly presented and that happy expressions were identified faster and with higher accuracy from faces than voices. Conversely, angry expressions were better recognized from voices than faces. Although no significant associations between single hormones (i.e., T or C) and RA or RTs were found, results showed that TxC interaction was significantly associated with both RA and RTs.

Our data highlights that the audio-visual presentation of emotional expressions significantly contributes to the ease and efficiency with which others’ emotions are recognized. This is in line with previous studies showing that the integration of auditorily and visually presented emotional information facilitates emotion recognition [e.g., Jessen et al., 2012; Paulmann & Pell, 2011; Baenziger et al., 2009), reflected in higher accuracy and faster RTs, especially for emotions such as disgust, fear (Collignon et al., 2008) and sadness (Kreifelts et al., 2007). One of the most noticeable differences between the present study and previous investigations was the presentation of several emotions and a neutral category (e.g., Collignon et al., 2008; De Gelder & Vroomen, 2000, included only two emotions) and the measurement of RTs (e.g., not considered in Kreifelts et al., 2007 study). Yet, the facilitation effect concerning stimulus classification manifested for every single emotion category during the audio-visual modality in comparison to the auditory modality. In addition, RA in the audio-visual modality exceeded that of the visual modality for angry, disgusted, neutral and sad emotions, which indicates the comprehensive nature of this integration process. As shown by the present results there are some differences in the effectiveness, with which specific emotions are recognized from voices and faces. Similar to the results reported in a meta-analysis by Elfenbein and Ambady (2002), anger was recognized better from voice than faces in our study, while better results for happiness were achieved from the visual compared to the auditory modality. This suggests that sensory modalities do not merely carry redundant information but rather, each may have certain specialized functions for the communication of emotions. Although the estimation of a visual threat (e.g., angry face) can be accurately predicted from close proximity, it has been shown that the louder, higher pitched sound of anger is particularly useful for both, proximal and distal spaces (see Ceravolo et al., 2016, for details). As it is highly adaptive to recognize and react to a potential threat in the environment (Pichon et al., 2008), the accurate detection of anger might, therefore, rely more on the human auditory than visual system. Previous research on facial expression recognition has consistently reported that happy expressions are recognized more accurately and faster than other basic emotions (e.g., Nummenmaa & Calvo, 2015). Our data provide further support for these findings, but not for our prediction (1b) that emotions communicated by the voice are recognized at higher rates of accuracy than in the visual channel. Nevertheless, it is possible that what determines the recognition advantage of happy faces is not so much their affect, but rather their perceptual and categorical distinctiveness from other emotional expressions (see Calvo et al., 2014, for details) as well as their frequent occurrence in everyday social contexts, thus, tuning the visual system towards efficient recognition of these faces (Nummenmaa & Calvo, 2015). Moreover, it has been argued that physical feature extraction can occur instantaneously for facial expressions, while the interplay of acoustic cues over time occurs in a probabilistic manner (Juslin & Laukka, 2003) and thus, may not engage a similar process for vocal expressions (see Paulmann & Pell, 2011, for details). This could have strengthened the underlying knowledge about emotions leading to improved RA and RTs in the visual modality.

The available evidence regarding the relationship between T and males’ emotion recognition ability is by no means clear-cut, making explicit claims about the direction of these effects impossible. The two predictions made in the present study were based on reported observations that T might have a negative influence on the recognition of emotions (Rukavina et al., 2018; Fujisawa & Shinohara, 2011), and that RTs of threat-related emotional expressions (i.e., angry, fear) would be much shorter with increasing levels of T (Derntl et al., 2009). To provide a more detailed picture of this association, we conducted an exploratory analysis for each modality and emotion category separately. In a similar fashion, we additionally analysed the effects of C. Similar to other reports in the literature, our data do not provide support for the influence of single steroid hormones (i.e., T or C) on RA or RTs (Duesenberg et al., 2016; Derntl et al., 2009). In contrast to the reported effect sizes or the significant effects between T and specific emotion categories (Rukavina et al., 2018; Derntl et al., 2009), the correlation coefficients for both hormones were small or close to zero across all modalities in our study. Despite our comparatively large sample, single hormones (i.e., T, C) did not appear to have an impact on RA and RTs in explicit emotion recognition tasks.

One assumption that has been put forth is that T and C do not act in isolation but rather interact to modulate complex social behaviours (Carré & Mehta, 2011). Following the dual-hormone hypothesis (Mehta & Josephs, 2010), we further explored whether the relationship between T and our response variables (i.e., RA and RT) is enhanced when C levels are low and attenuated when C levels are high. Similar to the obtained results in Dekkers et al. meta-analysis (2019) the overall effect size of T by cortisol interaction on RA and RT was significant but small in our study. Although our data support the dual-hormone hypothesis to some extent, they also showed that the interplay between T and C with RA or RTs is not as straightforward as one would expect. For instance, accuracy increased and RTs were shorter not only when T was high and C was low or vice-versa, but also when T and C were low. As our study is the first to account for the interaction between T and C on RA or RT, we cannot clearly provide explanations that might account for the observed mixed-pattern of results. However, as previous research found that high T and stress (C) levels impair cognitive abilities (e.g., Haenggi, 2004; Gouchie & Kimura, 1991) and decrease performance [e.g., Dolcos et al., 2014; Mehta et al., 2009), one would expect that with low levels of T and C, or with optimal levels of stress (i.e., eustress) but low T levels RA would increase in cognitive tasks. Since the pattern of the TxC interaction we found is unexpected and the effect size is small, we cannot rule out that it is a false-positive finding. Certainly, more work is needed to replicate our findings and to test these claims.

While our knowledge of how emotional information is integrated and recognized across channels is advancing steadily, the available literature, including the present study, is limited in a number of ways. In comparison to our study, most of the research mentioned above has evaluated a very small number of emotions (sometimes as few as two) and did not include a neutral baseline. Further, in some studies the audio material consisted of speech prosody (words, sentences). This opens up the possibility that the emotional tone of voice interacted with the affective value carried by the sentence’s/word’s semantic content. A related issue of past work is the use of emotional exemplars in conflict situations argued to be highly atypical of natural expressions of emotions (Paulmann & Pell, 2011). We addressed these issues by presenting emotion stimuli devoid of meaning (i.e., pseudo-words, pseudo-sentences and affect bursts) which always contained a congruent set of cues (i.e., encoder sex, stimulus time length) to express one of five basic emotions or a neutral state. We chose static faces to ensure our experimental conditions of stimulus presentation were compatible with the majority of prior literature. However, this format has been argued to be less ecologically valid (Krumhuber et al., 2013; Recio et al., 2011). While this assumption is still subject to some controversy (see Dobs et al., 2018, for details), future studies would benefit from using datasets of more naturalistic stimuli to further increase ecological validity.

As most of the previous research has focused on the associations between single hormones and facial emotion recognition, the present study uniquely contributes to the literature by providing a systematic examination of the influence of T, C and their interaction on RA and RT across different sensory modalities (i.e., auditory, visual and audio-visual). Although for C as well as for the interaction between T and C, the analyses were exploratory, they might prove of importance for researchers conducting work in this area to gain a more comprehensive understanding of when these effects emerge and when they do not. They may also yield a substantial theoretical payoff by enabling richer and more accurate predictions concerning the kind of outcomes tied to certain hormone level combinations.

The homogeneous characteristics of our sample (i.e., university students, narrow age range) may show patterns which do not hold for different sociodemographic subgroups. Given the increased focus on study replicability, future studies would benefit from combining datasets of different laboratories with similar outcome measures in order to reduce costs and increase the external validity, reliability and generalizability of findings. The present study provided evidence for differences in both RA and RTs in the three conditions of stimulus presentation and potentially set the stage regarding the influence of TxC interaction on these two response variables. It would thus be worthwhile to expand on these findings and examine whether the same holds true for the other sex. This could be done, for instance, by investigating the interaction between oestradiol and cortisol with RA, as previous research showed that high oestradiol is associated with more externalizing behaviours (linked to emotion-recognition difficulties, see Chronaki et al., 2015), but only when cortisol was low (Tackett et al., 2015).

## Conclusion

The findings of this study inform our current understanding with regard to the audio-visual integration of emotional signals among men by showing that audio-visual stimuli benefit RA over unimodal stimuli. They also explain inconsistencies in the past literature by highlighting that in explicit emotion recognition tasks voice-only expressions do not increase RA. Moreover, they replicate previous findings by establishing that for particular emotion categories RA and RTs vary as a function of modality. Crucially, our study contributes to a scientific domain that is currently reconsidering our understanding of the role hormones play for the recognition of emotions. It hereby paves the way for impactful future research, especially for the effects regarding TxC interaction with RA and RT.

## Acknowledgments

The authors thank Edmund Henniges for technical support, Saskia Brueckner, Marc Koehler and Isabel Noethen for help with data acquisition and all individuals who participated in the research presented here.

## Author contributions

A. L. designed the research with input from C. B., L. P. and A.S.; C. B. collected part of the data and wrote on the method part; A.L. analysed the data and wrote the paper with input from C. B., L. P. and A.S.

## Data availability

The dataset generated and analysed for the current study is available at osf.io/2ayms.

## Footnotes

1 All hypotheses tested in the current paper have been pre-registered (osf.io/w2tgr). This pre-registration contained further hypotheses that are not part of the present paper.

2 The data reported in this paper was obtained within the confines of a larger study. The experiment began with a short demographic questionnaire followed by the *Screening Questionnaire* (Schultheiss & Stanton, 2009), *Multi-Motive Grid* [MMG, Sokolowski et al., 2000] and *Positive and Negative Affect Schedule* [PANAS, Breyer & Bluemke, 2016]. Next, the first saliva sample (T1) was taken. After a short break, the ***Emotion Recognition Task*** ensued, followed by PANAS, and the collection of the second saliva sample (T2). The saliva samples were collected approximately 10 minutes before and after the emotion recognition task. The experiment ended with the completion of *Multifaceted Empathy Test* short-form [MET, Dziobek et al., 2008] and *Big Five Inventory* [BFI, Danner et al., 2016]. As MMG, PANAS, MET and BFI are not relevant to the present manuscript they are not further reported.

## References

Banse, R., Scherer, K. R., 1996. Acoustic profiles in vocal emotion expression. J. Pers. Soc. Psychol. 70, 614–636. http://dx.doi.org/10.1037/0022-3514.70.3.614

Belin, P., Fillion-Bilodeau, S., Gosselin, F., 2008. The Montreal affective voices: a validated set of nonverbal affect bursts for research on auditory affective processing. Behav. Res. Methods 40, 531–539. http://dx.doi.org/10.3758/BRM.40.2.531

Baenziger, T., Grandjean, D., Scherer, K., 2009. Emotion recognition from expressions in face, voice, and body: the multimodal emotion recognition test (MERT). Emotion 9, 691–704. https://doi.org/10.1037/a0017088

Breyer, B. Bluemke, M., 2016. Deutsche Version der Positive and Negative Affect Schedule PANAS (GESIS panel), in: Zusammenstellung Sozialwissenschaftlicher Items und Skalen. https://doi.org/10.6102/zis242

Calvo, M. G., García, A. G., Martín, A. F., Nummenmaa, L., 2014. Recognition of facial expressions of emotion is related to their frequency in everyday life. J. Nonverbal Behav. 38, 549–597. https://doi.org/10.1007/s10919-014-0191-3

Carré, J.M., Mehta, P. H., 2011. Importance of considering testosterone–cortisol interactions in predicting human aggression and dominance. Aggress. Behav. 37, 489–491. https://doi.org/10.1002/ab.20407

Ceravolo, L., Frühholz, S., Grandjean, D., 2016. Proximal vocal threat recruits the right voice-sensitive auditory cortex. Soc. Cogn. Affect. Neurosci. 11, 793–802. https://doi.org/10.1093/scan/nsw004

Chaplin, T. M., 2015. Gender and emotion expression: a developmental contextual perspective. Emot. Rev. 7, 14–21. https://doi.org/10.1177/1754073914544408

Chronaki, G., Wigelsworth, M., Pell, M.D., Kotz, S.A., 2018. The development of cross-cultural recognition of vocal emotions during childhood and adolescence. Sci. Rep. 8, 8659. https://doi.org/10.1038/s41598-018-26889-1

Chronaki, G., Garner, M., Hadwin, J.A., Thompson, M. J. J., Chin, C. Y., Sonuga-Barke, E. J. S., 2013. Emotion-recognition abilities and behavior problem dimensions in preschoolers: evidence for a specific role for childhood hyperactivity. Child Neuropsychol. 21, 25–40. https://doi.org/10.1080/09297049.2013.863273

Collignon, O., Girard, S., Gosselin, F., Roy, S., Saint-Amour, D., Lassonde, M., Lepore, F., 2008. Audio-visual integration of emotion expression. Brain Res. 1242, 126–135. https://doi.org/10.1016/j.brainres.2008.04.023

Cornew, L., Carver, L., Love, T., 2009. There’s more to emotion than meets the eye: a processing bias for neutral content in the domain of emotional prosody. Cogn. Emot. 24, 1133–1152. https://doi.org/10.1080/02699930903247492

Danner, D. et al. (2016). Die deutsche version des Big Five Inventory 2 (BFI-2), in Zusammenstellung sozialwissenschaftlicher items und skalen. https://doi.org/10.6102/zis247

Dekkers, T. J., Agelink van Rentergem, J. A., Meijera, B., Popmab, A., Wagemakera, E., Huizengaa, H. M., 2019. A meta-analytical evaluation of the dual-hormone hypothesis: does cortisol moderate the relationship between testosterone and status, dominance, risk taking, aggression, and psychopathy? Neurosci. Biobehav. Rev. 96, 250–271. https://doi.org/10.1016/j.neubiorev.2018.12.004

Derntl, B., Windischberger, C., Robinson, S., Kryspin-Exner, I., Gur, R. C., Moser, E., Habel, U., 2009. Amygdala activity to fear and anger in healthy young males is associated with testosterone. Psychoneuroendocrinology 34, 687–693. https://doi.org/10.1016/j.psyneuen.2008.11.007

Dobs, K., Bülthoff, I., Schultz, J., 2018. Use and usefulness of dynamic face stimuli for face perception studies – a review of behavioral findings and methodology. Front. Psychol. 9, 1355. https://doi.org/10.3389/fpsyg.2018.01355

Dolcos, F., Wang, L, Mather, M. 2014. Current research and emerging directions in emotion-cognition interactions. Front. Integr. Neurosci. 8, 83. https://doi.org/10.3389/fnint.2014.00083

Duesenberg, M., Weber, J., Schulze, L., Schaeuffele, C., Roepke, S., Regen, J. H., Otte, C., Wingenfeld, K., 2016. Does cortisol modulate emotion recognition and empathy? Psychoneuroendocrinology 66, 221–227. https://doi.org/10.1016/j.psyneuen.2016.01.011

Dziobek, I., Rogers, K., Fleck, S., Bahnemann, M., Heekeren, H. R., Wolf, O. T., Convit, A., 2008. Dissociation of cognitive and emotional empathy in adults with Asperger syndrome using the multifaceted empathy test (MET). J. Autism Dev. Disord. 38, 464–473. https://doi.org/10.1007/s10803-007-0486-x

de Gelder, B., Vroomen, J. 2000. The perception of emotions by ear and by eye. Cogn. Emot. 14, 289–311. https://doi.org/10.1080/026999300378824

Elfenbein, H. A., Ambady, N. 2002. On the universality and cultural specificity of emotion recognition: a meta-analysis. Psychol. Bull. 128, 203–235. https://doi.org/10.1037/0033-2909.128.2.203

Fischer, A. H., LaFrance, M. 2015. What drives the smile and the tear: why women are more emotionally expressive than men. Emot. Rev. 7, 22–29. https://doi.org/10.1177/1754073914544406

Fischer, A. H., Manstead, A. S. R., 2008. Social functions of emotion, in Lewis, M., Haviland-Jones, J., Barrett L. F. (Eds.), Handbook of emotions (3^rd^ Ed). Guilford Press, New York, pp. 456–468.

Fujisawa, T. X., Shinohara, K., 2011. Sex differences in the recognition of emotional prosody in late childhood and adolescence. J. Physiol. Sci. 61, 429–435. https://doi.org/10.1007/s12576-011-0156-9

Gignell, M., Hornung, J., Derntl, B., 2019. Emotional processing and sex hormones, in Schultheiss O. C., Mehta P. H. (Eds.), Routledge International Handbook of Social Neuroendocrinology (1^st^ Ed.). Routledge, Abingdon, UK, pp. 403–419. https://www.routledgehandbooks.com/doi/10.4324/9781315200439-24

Gouchie, C., Kimura, D., 1991. The relationship between testosterone levels and cognitive ability patterns. Psychoneuroendocrinology, 16, 323–334. https://doi.org/10.1016/0306-4530(91)90018-O

Grebe, N.M., Del Giudice, M., Thompson, M.E., Nickels, N., Ponzi, D., Zilioli, S., Mastripieri, D., Gangestad, S.W., 2019. Testosterone, Cortisol, and Status-Striving Personality Features: A Review and Empirical Evaluation of the Dual Hormone Hypothesis. Horm. Behav. 109, 25–37. https://doi.org/10.1016/j.yhbeh.2019.01.006

Hall, J. A., Carter, J. D., Horgan, T. G., 2000. Gender differences in the nonverbal communication of emotion, in Fischer A. H. (Ed.), Gender and Emotion: Social Psychological Perspectives, Cambridge University Press, Cambridge, pp. 97–117.

Hall, J. A., 1984. Non-Verbal Sex Differences. Communication Accuracy and Expressive Style. John Hopkins Press, London.

Haenggi Y., 2004. Stress and Emotion Recognition: An Internet Experiment Using Stress Induction. Swiss Journal of Psychology 63, 113–125. https://doi.org/10.1024/1421-0185.63.2.113

Hyde, J. S., 2014. Gender similarities and differences. Annu. Rev. Psychol. 65, 373–398. https://doi.org/10.1146/annurev-psych-010213-115057

Jessen, S., Obleser, J., Kotz, S. A., 2012. How bodies and voices interact in early emotion perception. PLoS ONE 7, e36070. http://dx.doi.org/10.1371/journal.pone.0036070

Juslin, P. N., Laukka, P., 2003. Communication of emotions in vocal expression and music performance: different channels, same code? Psychol. Bull. 129, 770–814. http://dx.doi.org/10.1037/0033-2909.129.5.770

Keltner, D., Kring, A.M., 1998. Emotion, social function and psychopathology. Rev. General Psychol. 2, 320–342. http://dx.doi.org/10.1037/1089-2680.2.3.320

Klasen, M., Kreifelts, B., Chen, Y. H., Seubert, J., Mathiak, K., 2014. Neural processing of emotion in multimodal settings. Front. Hum. Neurosci. 8, 822 http://dx.doi.org/10.3389/fnhum.2014.00822

Kordsmeyer, T. L., Lohöfener, M., Penke, L., 2018. Male facial attractiveness, dominance, and health and the interaction between cortisol and testosterone. Adapt. Human Behav. Physiol., 1–12. https://doi.org/10.1007/s40750-018-0098-z

Kosonognov, V., Titova, A., 2018. Recognition of all basic emotions varies in accuracy and reaction time: A new verbal method of measurement. Int. J. Psychol. 16, http://dx.doi.org/10.1002/ijop.12512

Krumhuber, E. G., Kappas, A., Manstead, A.S.R., 2013. Effects of dynamic aspects of facial expressions: a review. Emot. Rev. 5, 41–46. http://dx.doi.org/10.1177/1754073912451349

Lambrecht, L., Kreifelts, B., Wildgruber, D (2014). Gender differences in emotion recognition: impact of sensory modality and emotional category. Cogn. Emot. 28, 452–469. http://dx.doi.org/10.1080/02699931.2013.837378

Langner, O., Dotsch, R., Bijlstra, G., Wigboldus, D. H. J., Hawk, S. T., van Knippenberg, Ad, 2010. Presentation and validation of the Radboud Faces Database. Cogn. Emot. 24, 1377–1388. http://dx.doi.org/10.1080/02699930903485076

Kraus, M. W., 2017. Voice-only communication enhances empathic accuracy. Am. Psychol. 72, 644–654. http://dx.doi.org/10.1037/amp0000147

Kreifelts, B., Ethofer, T., Grodd, W., Erb, M., Wildgruber, D., 2007. Audiovisual integration of emotional signals in voice and face: An event-related fMRI study. NeuroImage 37, 1445–1456. https://doi.org/10.1016/j.neuroimage.2007.06.020.

Lausen, A., Hammerschmidt, K., Schacht, A., 2019. Emotion recognition and confidence ratings predicted by vocal stimulus type and acoustic parameters. https://doi.org/10.31234/osf.io/kqy2n

Lausen, A., Schacht, A., 2018. Gender differences in the recognition of vocal emotions. Front. Psychol. 9, 882. https://doi.org/10.3389/fpsyg.2018.00882

Mazur, A., Booth, A., 2014. Testosterone is related to deviance in male army veterans, but relationships are not moderated by cortisol. Biol. Psychol. 96, 72–76. https://doi.org/10.1016/j.biopsycho.2013.11.015

Mehta, P.H., Prasad, S., 2015. The dual-hormone hypothesis: A brief review and future research agenda. Curr. Opin. Behav. Sci. 3, 163–168. https://doi.org/10.1016/j.cobeha.2015.04.008

Mehta, P.H., Josephs, R.A., 2010. Testosterone and cortisol jointly regulate dominance: Evidence for a dual-hormone hypothesis. Horm. Behav. 58, 898–906. https://doi.org/10.1016/j.yhbeh.2010.08.020

Mehta, P.H., Wuehrmann, E.V. Josephs R. A. 2009. When are low testosterone levels advantageous? The moderating role of individual versus intergroup competition. Horm. Behav. 56, 158–162. https://doi.org/10.1016/j.yhbeh.2009.04.001

Montagne, B., Kessels, R. P., de Haan, E. H. F., Perrett, D. I., 2007. The emotion recognition task: a paradigm to measure the perception of facial emotional expressions at different intensities. Percept. Mot. Skills 104, 589–598. https://doi.org/10.2466/pms.104.2.589-598

Nummenmaa, L., Calvo, M. G., 2015. Dissociation between recognition and detection advantage for facial expressions: a meta-analysis. Emotion 15, 243–256. http://dx.doi.org/10.1037/emo0000042

Palermo, R. Coltheart, M., 2004. Photographs of facial expression: accuracy, response times, and ratings of intensity. Behav. Res. Methods Instrum. Comput. 36, 634–638. https://doi.org/10.3758/BF03206544

Paulmann, S., Pell, M. D., 2011. Is there an advantage for recognizing multi-modal emotional stimuli? Motiv. Emot. 35, 192–201. https://doi.org/10.1007/s11031-011-9206-0

Paulmann, S., Kotz, S.A., 2008. An ERP investigation on the temporal dynamics of emotional prosody and emotional semantics in pseudo- and lexical sentence context. Brain Lang. 105, 59–69. http://dx.doi.org/10.1016/j.bandl.2007.11.005

Paulmann, S., Pell, M.D., Kotz, S.A., 2008. How aging affects the recognition of emotional speech, Brain Lang. 104, 262–269. http://dx.doi.org/10.1016/j.bandl.2007.03.002

Pell, M.D., Rothermich, K., Liu, P., Paulmann, S., Sethi, S., Rigoulot, S., 2015. Preferential decoding of emotion from human non-linguistic vocalizations versus speech prosody. Biol. Psychol. 111, 14–25. http://doi.org/10.1016/j.biopsycho.2015.08.008

Pichon, S., de Gelder, B., Grèzes, J., 2008. Emotional modulation of visual and motor areas by dynamic body expressions of anger. Soc. Neurosci. 3, 199–212. http://dx.doi.org/10.1080/17470910701394368

Piwek, L., Pollick, F., Petrini, K., 2015. Audiovisual integration of emotional signals from others’ social interactions. Front. Psychol. 6, 611. http://dx.doi.org/10.3389/fpsyg.2015.00611

R Core Team, 2017. R: A Language and Environment for Statistical Computing. Vienna: R Foundation for Statistical Computing. Available online at: https://www.r-project.org

Recio, G., Sommer, W., Schacht, A., 2011. Electrophysiological correlates of perceiving and evaluating static and dynamic facial emotional expressions. Brain Res. 1376, 66–75. https://doi.org/10.1016/j.brainres.2010.12.041

Rukavina, S., Sachsenweger, F., Jerg-Bretzke, L., Daucher, A. E., Traue, H. C., Walter, S., Hoffmann, H., 2018. Testosterone and its influence on emotion recognition in young, healthy males. Psychology 9, 1814–1827. http://dx.doi.org/10.4236/psych.2018.97106

Sarkar, A., Mehta, P. H., Josephs, R. A., 2019. The dual-hormone approach to dominance and status-seeking, in: Schultheiss O. C., Mehta P. H. (Eds.), Routledge International Handbook of Social Neuroendocrinology (1^st^ Ed.), Routledge, Abingdon, UK, pp. 113–132. https://www.routledgehandbooks.com/doi/10.4324/9781315200439-8

Schirmer, A., 2013. Sex differences in emotion, in: Armony, J., Vuilleumier, P. (Eds.), The Cambridge Handbook of Human Affective Neuroscience (1^st^ Ed.). Cambridge University Press, New York, pp. 591–611.

Schultheiss, O. C., Dlugash, G., Mehta, P. H., 2019. Hormone measurement in social neuroendocrinology: a comparison of immunoassay and mass spectrometry methods, in: Schultheiss O. C., Mehta P. H. (Eds.), Routledge International Handbook of Social Neuroendocrinology (1^st^ Ed.), Routledge, Abingdon, UK, pp. 26–41. https://www.routledgehandbooks.com/doi/10.4324/9781315200439-3

Schultheiss, O. C., Schiepe, A., Rawolle, M., 2012. Hormone assays, in: Cooper, H., Camic, P. M., Long, D. L., Panter, A. T., Rindskopf, D., Sher, K. J. (Eds.), Handbook of research methods in psychology (Vol. 1), American Psychological Association, Washington, pp. 489–500.

Schultheiss, O. C., Stanton, S. J., 2009, in: Harmon-Jones, E. & Beer, J. S. (Eds.), Methods in social neuroscience, Guilford Press: New York, pp. 17–44.

Sokolowski, K., Schmalt, H. D., Langens, T. A., Puca, R. M., 2000. Assessing achievement, affiliation, and power motives all at once: The Multi-Motive Grid (MMG). J. Pers. Assess. 74, 126–145.

Soto, J.A., Levenson, R.W., 2009. Emotion recognition across cultures: the influence of ethnicity on empathic accuracy and physiological linkage. Emotion 9, 874–884. http://dx.doi.org/10.1037/a0017399

Tackett, J. L., Reardon, K. W., Herzhoff, K., Page-Gould, E., Harden, K. P., Josephs, R.A., 2015. Estradiol and cortisol interactions in youth externalizing psychopathology. Psychoneuroendocrinology 55, 146–153. http://dx.doi.org/10.1016/j.psyneuen.2015.02.014

Thompson, A. E., Voyer, D., 2014. Sex differences in the ability to recognize non-verbal displays of emotion: A meta-analysis. Cogn. Emot. 28, 1164–1195. http://dx.doi.org/10.1080/02699931.2013.875889

Viau, V., 2002. Functional cross-talk between the hypothalamic-pituitary-gonadal and- adrenal axes. J. Neuroendocrinol., 14, 506–513. http://dx.doi.org/10.1046/j.1365-2826.2002.00798.x

Vongas, J. G., Hajj, R. A., 2017. The effects of competition and implicit power motive on men’s testosterone, emotion recognition, and aggression. Horm. Behav. 92, 57–71. http://dx.doi.org/10.1016/j.yhbeh.2017.04.005

Waaramaa, T., 2017. Gender differences in identifying emotions from auditory and visual stimuli. Logop. Phoniatr. Voco. 42, 160–166. https://doi.org/10.1080/14015439.2016.1243725

Wagner, H. L., 1993. On measuring performance in category judgement studies of nonverbal behaviour. J. Nonverbal Behav. 17, 3–28. http://dx.doi.org/10.1007/BF00987006

Wells L. J., Gillespie S. M., Rotshtein, P., 2016. Identification of emotional facial expressions: effects of expression, intensity, and sex on eye Gaze. PLoS ONE 11, e0168307. https://doi.org/10.1371/journal.pone.0168307

Welker, K.M., Lozoya, E., Campbell, J.A., Neumann, C.S., Carré, J.M., 2014. Testosterone, cortisol, and psychopathic traits in men and women. Physiol. Behav. 129, 230–236. https://doi.org/10.1016/j.physbeh.2014.02.057

Wendt, B., Scheich, H., 2002. The “Magdeburger Prosodie Korpus” – a spoken language corpus for fMRI-Studies, in: Bel, B., Marlien, I. (Eds.), Speech Prosody (pp.699–701). Aix-en-Provence: SproSIG

Williams, L. M., Mathersul, D., Palmer, D. M., Gur, R. C., Gur, R. E., Gordon, E., 2009. Explicit identification and implicit recognition of facial emotions: I. Age effects in males and females across 10 decades. J. Clin. Exp. Neuropsychol. 31, 257–277. http://dx.doi.org/10.1080/13803390802255635

World Medical Association (2013). World Medical Association Declaration of Helsinki: ethical principles form medical research involving human subjects. JAMA 310, 2191–2194. http://dx.doi.org/10.1001/jama.2013.281053

